# Characterizing the spatial signal of environmental DNA in river systems using a community ecology approach

**DOI:** 10.1101/2020.10.11.333047

**Authors:** Isabel Cantera, Jean-Baptiste Decotte, Tony Dejean, Jérôme Murienne, Régis Vigouroux, Alice Valentini, Sébastien Brosse

**Affiliations:** Laboratoire Evolution et Diversité Biologique, UMR5174, CNRS, IRD, Université Toulouse III Paul Sabatier, 118 route de Narbonne, F-31062 Toulouse, France.; VIGILIFE, 17 rue du Lac Saint-André Savoie Technolac - BP 274, Le Bourget-du-Lac 73375, France.; SPYGEN, 17 rue du Lac Saint-André Savoie Technolac - BP 274, Le Bourget-du-Lac 73375, France.; HYDRECO, Laboratoire Environnement de Petit Saut, B.P 823, F-97388, Kourou Cedex, French Guiana.

**Keywords:** Neotropical fishes, eDNA, spatial signal, distance decay, distribution range, freshwater

## Abstract

Environmental DNA (eDNA) is gaining a growing popularity among scientists but its applicability to biodiversity research and management remains limited in river systems by the lack of knowledge about the spatial extent of the downstream transport of eDNA.

Here, we assessed the ability of eDNA inventories to retrieve spatial patterns of fish assemblages along two large and species rich Neotropical rivers. We first examined overall community variation with distance through the distance decay of similarity and compared this pattern to capture-based samples. We then considered previous knowledge on individual species distributions, and compared it to the eDNA inventories for a set of 53 species.

eDNA collected from 28 sites in the Maroni and 25 sites in the Oyapock rivers permitted to retrieve a decline of species similarity with distance between sites. The distance decay of similarity derived from eDNA was similar, and even more pronounced, than that obtained with capture-based methods (gil-nets). In addition, the species upstream-downstream distribution range derived from eDNA matched to the known distribution of most species.

Our results demonstrate that environmental DNA does not represent an integrative measure of biodiversity across the whole upstream river basin but provide a relevant picture of local fish assemblages. Importantly, the spatial signal gathered from eDNA was therefore comparable to that gathered with local capture based methods, which describes fish fauna over a few hundred metres.

## Introduction

In aquatic ecosystems, DNA molecules flowing in the water after being separated from their source organisms can be assigned to taxa using the environmental DNA (eDNA) metabarcoding approach. The method consists on the sampling, extraction, amplification and sequencing of free DNA suspended in the water or linked to suspended particles. Subsequently, the obtained sequences are taxonomically assigned to species based on comparisons to a reference molecular database (Taberlet, Bonin, Zinger, & Coissac, 2018). This approach has proven to be particularly efficient at measuring the diversity of fishes in aquatic ecosystems (Civade et al., 2016; Hänfling et al., 2016; Olds et al., 2016; Valentini et al., 2016; Evans et al., 2017; Pont et al., 2018; Cantera et al., 2019). Indeed, in those studies, the eDNA samples provided similar or more complete inventories than traditional survey methods, such as netting, electrofishing or visual records, at the same sites. The great detection performance of the method may be due to its high sensitivity to detect species that are rare and difficult to sample with traditional methods (Ficetola, Miaud, Pompanon, & Taberlet, 2008; Goldberg, Pilliod, Arkle, & Waits, 2011; Jerde, Mahon, Chadderton, & Lodge, 2011). In fact, the species list obtained with one eDNA sampling session was often comparable to cumulative traditional sampling sessions and historical records (Civade *et al*. 2016; Hänfling *et al*. 2016; Nakagawa *et al*. 2018; Pont *et al*. 2018; Cantera et al. 2019). Due to its sampling efficiency to reconstruct whole aquatic communities, eDNA methods are expected to revolutionize survey methods for ecological research and biodiversity monitoring on aquatic ecosystems (Keck, Vasselon, Tapolczai, Rimet, & Bouchez, 2017; Rees, Gough, Middleditch, Patmore, & Maddison, 2015; Rees, Maddison, Middleditch, Patmore, & Gough, 2014; Zinger et al., 2020). This is particularly true for large rivers where sampling the fauna can be very difficult, as it is time consuming, costly and limited to specific habitats (Pont et al., 2018).

The application of eDNA has been however limited in rivers because the species detection recovered by eDNA sampling might be decoupled from the species’ physical habitat due to downstream eDNA transport. The network structure of rivers is governed by a continuous and directional flow (McCluney et al., 2014) that can transport eDNA downstream from the DNA source while it degrades (Barnes & Turner, 2016). The spatial extent of downstream transport depends on the degradation rate of eDNA (Barnes & Turner, 2016) combined with the discharge rates of the water system (Jane et al., 2015). For instance, fast-flow systems may transport eDNA downstream over large distances before its degradation and dilution. Moreover, the identity and density of target species could also influence eDNA detection distance (Deiner & Altermatt, 2014; Pilliod, Goldberg, Arkle, & Waits, 2014; Pont et al., 2018).

In aquaria and mesocosms, eDNA remained detectable from a few days to a few weeks after its release in the water (see Table 1, Barnes and Turner 2016). Besides being heterogeneous, these experimental results are difficult to relate with water velocity in the field to estimate eDNA detection distance, given that natural ecosystems are more complex and local environmental conditions, which influence eDNA degradation rate (Barnes & Turner, 2016), are covarying. For instance, Takahara *et al*. (2012) found a significant relationship between eDNA production from common carp and water temperature in a natural lagoon but not in aquaria. Moreover, in experimental designs the DNA of target species are usually in higher concentrations compared to natural ecosystems while DNA concentration was shown to influence degradation rate (Barnes & Turner, 2016).

Deiner *et al*. (2016) advocated that eDNA-based inventories represent a spatially integrated measure of biodiversity describing the diversity at the catchment scale and considered that rivers act as conveyor belts for eDNA. Therefore, eDNA collected at one sampling point may represent the combination of local and upstream diversity from this point. Indeed, the assumption that eDNA of entire communities will accumulate downstream omits the phenomenon of DNA degradation and sedimentation once it has been shed from an organism (Barnes and Turner 2016; Pont et al. 2018). The detection distance of eDNA was investigated more explicitly in natural rivers and streams in temperate ecosystems. Although the studies agreed that eDNA from a species can be detected downstream from where the species occurs and that eDNA signal decays over distance, the estimation of detection distance displays a great variability among studies. The introduction of caged animals in streams revealed that eDNA was detected 5 m but not 50 m downstream from caged Idaho giant salamander (Pilliod et al., 2014) and up to 1 km downstream from caged brook trout (Wilcox et al., 2016). Contrastingly, eDNA from lake species has been found in downstream streams over greater distances. DNA from a lacustrine fish species was detected up to 3 km downstream from the outlet of a lake by Civade *et al*. (2016), and Deiner and Altermatt (2014) detected two lake invertebrate species (a planktonic crustacean and a mollusc) up to 12.3 km downstream from a lake. Nakagawa *et al*. (2018) assessed detection distance by comparing eDNA fish inventories with observational data within intervals from 1 to 10 km upstream from the eDNA sampling site, and showed eDNA inventories were more similar with observational data when comparing with traditional inventories within 6 km upstream from the eDNA sampling site. Pont *et al*. (2018) detected eDNA of Whitefish, an abundant species in Lake Geneva, up to 130 km downstream from the lake, whereas the detection of less abundant species was restricted to a few kilometres, suggesting that abundant species are detected farther downstream than rare ones.

The variability among detection distances may reflect the specific local environmental conditions of the studied water bodies (Barnes & Turner, 2016; Jane et al., 2015). Local conditions such as acidity, oxygen demand, primary production, water temperature, solar radiation, organic materials and the activity of microorganisms, have been shown to influence eDNA degradation rate (Barnes & Turner, 2016; Eichmiller, Best, & Sorensen, 2016; Seymour et al., 2018). As all the above-mentioned characteristics as well as discharge rate vary strongly between rivers, the spatial extent of downstream transport of eDNA may be highly dependent of the studied system. For example, by treating eDNA downstream transport as the vertical transfer of fine particulate organic matter from the water column to the riverbed, Pont *et al*. (2018) suggested that detection distance varies with river size, ranging from a few km in small streams to more than 100 km in large rivers.

Overall, the discrepancy of detection distances among species and rivers reported in the literature makes difficult to know whether the species recorded in an eDNA sample reflect a local fish assemblage or a more spatially extended synthesis of the species present in the whole river basin located upstream from eDNA sampling sites. Such uncertainty is reinforced in tropical environments, which host rich, but understudied, fish communities. Tropical rivers are also characterised by distinct environmental conditions than temperate and cold river ecosystems, where most developments of eDNA method were achieved (Zinger *et al*. 2020). Moreover, given the deep human-induced changes of freshwater fish biodiversity throughout the world (Su et al., 2021), it is now urgent to determine if the fast and non-destructive eDNA samples provide similar biodiversity measures to the traditional time consuming and often destructive capture-based methods.

Here, we aimed at characterizing the spatial signal of eDNA in two large Neotropical rivers through two complementary approaches considering: i) spatial patterns of the whole fish community and ii) the individual spatial range of species along the upstream-downstream river continuum. Considering the whole fish community, we sought to describe the decay of similarity between communities with increasing distance between sampling sites. This widely recognised pattern in ecology postulates that the similarity between local measures of biodiversity should decline with the distance between observations (Soininen *et al*. 2007; Morlon *et al*. 2008, Nekola and White 1999). This pattern should be retrieved if eDNA detection distance is short, whereas it should be blurred under the hypothesis of a long-distance transport of eDNA. We test this hypothesis in the Maroni and Oyapock rivers, two species rich Neotropical rivers differing in their anthropization levels. We predict a steeper distance decay in the Maroni river due to an upstream-downstream increase of anthropic disturbances (Gallay *et al*. 2018), which co-varies with environmental changes along the course of the river. We also compared the eDNA distance decay pattern to capture-based data available from the Maroni river. We here hypothesise that under a short eDNA detection distance, distance decay of similarity between eDNA and capture based methods should be the same. Regarding each fish species, we compared the distribution range of detected species along the upstream-downstream gradient of the Maroni river to the distribution range recorded by capture-based data in the same river. Again, under the hypothesis of a local eDNA detection of the species, the distribution range of species retrieved using eDNA should fall within the known species range, while under the hypothesis of a long-distance transport, species should be detected downstream from their known distribution range.

## Materials and methods

### Study sites

This study was conducted in two tropical rivers located in the Northern East of the Amazonian region (*sensu lato*, including Guiana shield and Amazon river drainage) (Figure 1). The Maroni river measures 612 km from the source to the estuary and its drainage basin covers a surface of more than 68 000 km^2^ in Suriname and French Guiana. The Oyapock river (404 km long and 26 800 km^2^) flows over Amapa (Brazil) and French Guiana. Both rivers are characterized by a low topographic gradient (mean slope =0.4% and 0.5% for Maroni and Oyapock respectively), warm water (30.5 ± 0.23 °C and 28.4 ± 0.35 °C) and neutral pH (7 ± 0.05 and 7.2 ± 0.03). Both rivers host typical freshwater Amazonian fauna with more than 270 and 190 described fish species in the Maroni and Oyapock, respectively (Le Bail et al., 2012). Among these species, 141 and 119 strictly freshwater species (estuarine species were excluded) have been recorded in the main channel of the Maroni and Oyapock rivers, respectively (Planquette *et al*. 1996; Keith *et al*. 2000). Despite being located in one of the largest unfragmented rainforest areas in the world, the Maroni river is facing an unprecedented rise of land use changes due mainly to gold-mining activities but also to agriculture and urbanization (Gallay et al., 2018; Rham et al., 2017). Such anthropogenic disturbances show an increasing gradient along the course of the Maroni. Upstream areas are free of human settlements and part of a National Park, whereas human densities as well as deforestation and mining intensities are increasing toward the downstream part of the river (Gallay et al., 2018). Thus, environmental and anthropogenic gradients are covarying along the Maroni river. Contrastingly, the Oyapock river remains less impacted, with low human population density and gold mining activities limited to a few tributaries (Gallay et al., 2018).

**Figure 1:**
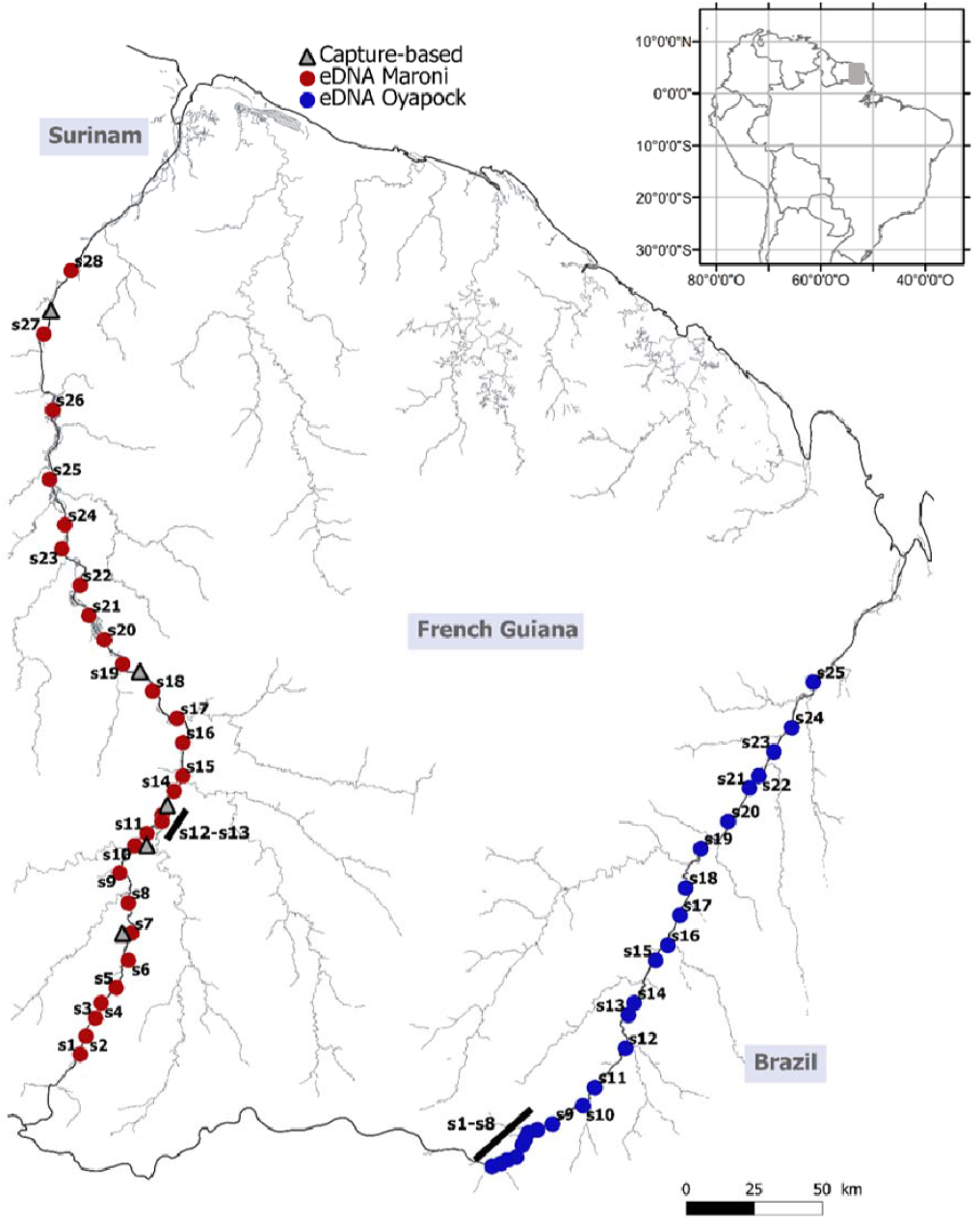
Map of the study area indicating the eDNA sampling sites in the Maroni (red dots) and Oyapock (blue dots) rivers. Grey triangles correspond to sites sampled with a capture-based method (gill-nets). Inset map on the top indicate the location of the study area in South America.

### eDNA sampling

We collected environmental DNA samples in 53 sites along the main channel of two rivers (Figure 1), with 28 sites sampled on the Maroni main channel and 25 sites sampled on the Oyapock main channel. The estuary sections, from the sea to the upstream area influenced by the tide, were not considered in this study, because tidal water movements could bias our results through downstream-upstream movement of eDNA during the rising tide. The sampling was achieved during a 2 weeks field session during the dry season (October-November) in 2017 and 2018 for the Maroni and Oyapock, respectively. In all sites, the river was wider than 20 meters and deeper than one meter (Strahler orders 4 to 8). The distance between adjacent sites ranged from 1.6 to 31.8 km (mean: 13.8 and SE: 1.4) in the Maroni, and from 1.2 to 23.8 km (mean: 10.8 and SE: 1.5) in the Oyapock. In both rivers, sites were sequentially sampled from downstream to upstream, with 1 to 4 sites sampled per day, according to the travel time between sites. All the metadata associated with the samples are available through the CEBA geoportal (vmcebagn-dev.ird.fr) or the French Guiana geoportal (geoguyane.fr) under reference 5617a9ff-d0aa-48a9-b2c2-cb7fd5b92692 (Murienne et al., 2019).

Following the protocol implemented by Cantera et al. (2019), we filtered two samples of 34 liters of water at each site to collect eDNA. A peristaltic pump (Vampire sampler, Burlke, Germany) and a single-use tubing were used to pump the water into a single-use filtration capsule (VigiDNA 0.45 μm; SPYGEN, le Bourget-du-Lac, France). All the material used to collect water was single use, and therefore replaced between each sample. Single use gloves were also used to avoid contamination. To collect eDNA, the input part of the tubing was placed 10 to 20 centimetres below the surface in areas with high water flow as recommended by Cilleros *et al*. (2019). Sampling was achieved in turbulent area (rapid hydromorphologic unit) to ensure an optimal homogenization of the eDNA throughout the water column. To avoid DNA contamination among sites, the operator always remained downstream from the filtration area and stayed on emerging rocks. At the end of the filtration, the filtration capsule was emptied of water, filled with 80 mL of CL1 conservation buffer (SPYGEN) and stored in individual sterile plastic bags kept in the dark.

### Capture-based sampling

The fish captures were performed using a standardized gill-net protocol designed by Tejerina-Garro and De Mérona (2001) and used as a routine biodiversity monitoring protocol by the French Ministry of Environment (DGTM), as part of the European Water Framework directive. Five sites located on the Maroni main channel were sampled yearly, during the dry season, from 2007 to 2016. Briefly, the protocol consist in sampling fish overnight using ten gill-nets measuring 25 metre long and 1.5 metre height and with mesh sizes ranging from 10 to 70 millimetres. The 10 nets are set before down, over a river reach measuring 300 to 500 metres. Nets were removed 12 hours later, after down, and all captured fish were identified to the species level. This protocol allows capturing both diurnal and nocturnal species, and most of the fish size ranges except the smallest species. Yearly data were pooled to provide a broad characterisation of the fish fauna in each site. This permitted to deal with the limited efficiency of gill-nets to collect the entire fauna. Indeed, gill-nets are strongly dependent of fish activity and need repeated sampling effort to provide realistic species community inventories (Murphy and Willis 1996; Pont *et al*. 2018).

### eDNA laboratory and bioinformatic analyses

For DNA extraction, each filtration capsule was agitated for 15 min on an S50 shaker (cat Ingenieurbüro™) at 800 rpm and then emptied into a 50 mL tube before being centrifuged for 15 min at 15,000×g. The supernatant was removed with a sterile pipette, leaving 15 mL of liquid at the bottom of the tube. Subsequently, 33 mL of ethanol and 1.5 mL of 3M sodium acetate were added to each 50 mL tube and stored for at least one night at −20°C. Tubes were centrifuged at 15 000 ×g for 15 min at 6°C, and the supernatants were discarded. After this step, 720 μL of ATL buffer from the DNeasy Blood & Tissue Extraction Kit (Qiagen) was added. The tubes were then vortexed, and the supernatants were transferred to 2 mL tubes containing 20 μL of proteinase K. The tubes were finally incubated at 56°C for two hours. Afterwards, DNA extraction was performed using NucleoSpin Soil (Macherey-Nagel GmbH & Co., Düren Germany) starting from step six and following the manufacturer’s instructions. The elution was performed by adding 100 μL of SE buffer twice. Four negative extraction controls were also performed. They were amplified and sequenced in parallel to the field samples to monitor possible laboratory contaminants. After the DNA extraction, the samples were tested for inhibition by qPCR (Biggs et al. (2015). If the sample was considered inhibited, it was diluted 5-fold before the amplification.

We performed DNA amplifications in a final volume of 25 μL including 1 U of AmpliTaq Gold DNA Polymerase (Applied Biosystems, Foster City, CA), 10 mM of Tris-HCl, 50 mM of KCl, 2.5 mM of MgCl_2_, 0.2 mM of each dNTP, 0.2 μM of “teleo” primers (Valentini et al., 2016) and 3 μL of DNA template. We also added human blocking primer for the “teleo” primers with a final concentration of 4 μM and 0.2 μg/μL of bovine serum albumin (BSA, Roche Diagnostic, Basel, Switzerland) to the mixture. We performed 12 PCR replicates per field sample. The forward and reverse primer tags were identical within each PCR replicate. The PCR mixture was denatured at 95°C for 10 min, followed by 50 cycles of 30 s at 95°C, 30 s at 55°C and 1 min at 72 °C and a final elongation step at 72°C for 7 min. This step was done in a room dedicated to amplified DNA with negative air pressure and physical separation from the DNA extraction rooms (with positive air pressure). We also amplified 14 negative extraction controls and five PCR negatives controls and sequenced them in parallel with the PCR replicates. We pooled the purified PCR products in equal volumes to achieve an expected sequencing depth of 500,000 reads per sample before the libraries preparation. Ten libraries were prepared using a PCR-free library protocol (https://www.fasteris.com/metafast), at Fasteris facilities (Geneva, Switzerland). Four libraries were sequenced using an Illumina HiSeq 2500 (2×125 bp) (Illumina, San Diego, CA, USA) and the HiSeq SBS Kit v4, three using a MiSeq (2×125 bp) (Illumina, San Diego, CA, USA) and the MiSeq Flow Cell Kit Version3 (Illumina, San Diego, CA, USA) and three using a NextSeq (2×150 bp+8bp) (Illumina, San Diego, CA, USA) and the NextSeq Mid kit (Illumina, San Diego, CA, USA). The libraries ran on the NextSeq were equally distributed in four lanes. Sequencing were performed following the manufacturer’s instructions at Fasteris facilities (Geneva, Switzerland).

The sequence reads were analyzed using the OBITools package (Boyer et al., 2016) following the protocol described in Valentini et al. (2016). The ecotag program was used for the taxonomic assignment of molecular operational taxonomic units (MOTUs). We kept only the MOTUS with a similarity to our reference database above 98%. Our local reference database represent an updated version of the reference database available in Cilleros *et al*. (2019), resulting in 251 Guianese fish species. The GenBank nucleotide database was checked but Guianese fishes being poorly referenced (most of the sequences are from Cilleros *et al*. (2019)), it did not provided additional information in our case. We discarded all MOTUs with a frequency of occurrence below 0.001 per library in each sample, considered as tag-jumps (Schnell, Bohmann, & Gilbert, 2015). These thresholds were empirically determined to clear all reads from the extraction and PCR negative controls included in our global data production procedure as suggested by De Barba et al. (2014) and Taberlet et al. (2018). For the samples sequenced on a NextSeq platform, only species present in at least two lanes were kept. The entire procedure was repeated for each sample, giving rise to two species lists per field site. The two obtained species lists per site were finally pooled to obtain a single species inventory per site, as Cantera *et al*. (2019) showed that pooling two replicate field samples was sufficient to inventory most of the fauna in highly diverse tropical aquatic ecosystems from the same region.

### eDNA data ability to reveal spatial patterns in ecology

We first tested the ability of eDNA data to reveal patterns of decay of similarity in species composition between pairs of sites with increasing geographic distance. Presence/absence matrices of the species detections in each site were used to build a species compositional distance matrix between pairs of sites, for each river separately with the *vegdist* function. Pairwise compositional distances between sites were calculated using the Jaccard index. We used linear regressions to model the relationship between species similarity distances and watercourse distances between sites (in km) for each river. Considering that the environment have been shown to influence the rate of similarity decay (Nekola & White, 1999), differences in the rates of decay in similarity with distance between rivers were tested. To do this, the slopes and intercepts of the obtained decay relationships were compared by using a permutation procedure with 999 iterations based on linear regressions. The permutation procedure was done with the *diffslope* and *diffic* functions of the ‘simba’ R package for the slope and intercept comparisons, respectively. This function compares the observed difference between the slopes or intercepts of each linear regression from each river with the permuted slope differences by randomly interchanging the values between the two river datasets. The p-value is computed as the ratio between the number of cases where the differences in slopes exceed the difference in slope of the actual configuration.

The distance decay pattern obtained with eDNA samples between the Oyapock and Maroni rivers was compared. Then, we did the same between eDNA and capture-based data from the Maroni river (Figure 1). The comparison between the eDNA and capture-based distance decay profiles was first made with all the species detected (or captured) by the two methods. To deal with differences in species capturability between methods, and particularly with the low capturability of small (less than 10 centimetres long) or non-mobile species using gill-nets. Finally, we restricted the comparison to the species detected (or captured) with both methods.

For the species level assessment, the spatial upstream-downstream range of each detected species was compared to the known distribution range of the species derived from the capture-based method. We used the watercourse distance from the river mouth (in km) of the sampled sites to represent the spatial distribution range along the upstream-downstream gradient. To keep the spatial range comparable between the two sampling methods, we discarded the most upstream sites sampled by eDNA (S1 to S6, see Figure 1), as this part of the Maroni river has not been investigated using gill nets. Moreover, gill-net being selective for species (*e.g*. small and/or elongated species are not captured) we restricted this analysis to the species detected by both eDNA and by gill-nets in at least two sites or sampling events. Indeed, the 11 species detected only once over the 9 years’ gill-net surveys represent “bycatches” of small species (<10 cm) such as *Characidium zebra* or *Hemigrammus surinamensis*, which are hardly captured using gill-nets. We here predict that under a short eDNA detection distance, species detections using eDNA should not systematically occur downstream from the known distribution range of the species as established by gill-net data.

## Results

Fish species were detected in all the eDNA replicates. The number of species per site ranged from 50 to 71 (mean = 62) in the Oyapock sites and 30 to 94 (mean = 57) in the Maroni sites. We detected a total of 160 species, with 116 and 96 species in the Oyapock and Maroni rivers, respectively. They account for more than 80% (80.7% and 82.3% for the Oyapock and Maroni, respectively) of the fish species known from the main channel of both river basins, excluding estuarine species (Le Bail, Keith, & Planquette, 2000; Planquette, Keith, P, & Le Bail, 1996). Among the five Maroni sites sampled with capture-based methods (gill-nets), a total of 94 fish species were captured, with species richness ranging from 50 to 66 per site. The comparison of eDNA (22 sites, sites S7 to S28, Figure 1) and gill-nets (5 sites) inventories revealed that 53 species were detected by both methods in the Maroni river.

Results from the linear models showed that similarity decreased significantly with watercourse distance between sites pairs sampled with eDNA data in both Oyapock (p<0.001, slope= −0.0011, R^2^=0.5) and Maroni (p<0.001, slope= −0.00125, R^2^=0.6) rivers. As sites became more distant, their corresponding fish communities became more dissimilar in species (Figure 2**A**). For both rivers, the similarity decay with distance relationship fitted well without obvious change of the slope of the regression in any distance value. The permutation procedure showed that the negative effect of spatial distance on species similarity was more marked in the Maroni than in the Oyapock river, with a significantly more negative slope (permutation test, p = 0.02) on the Maroni river.

**Figure 2:**
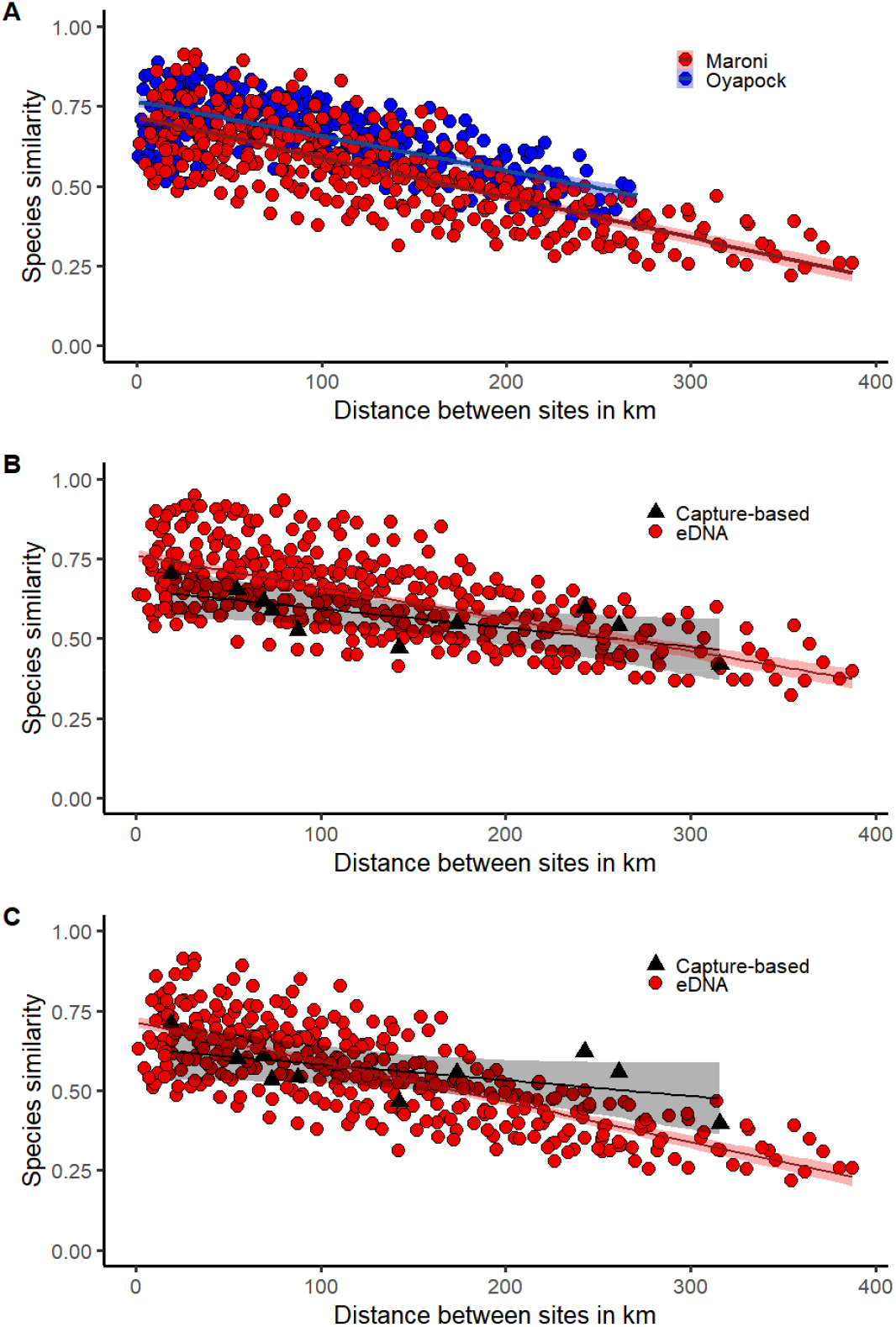
Decay of fish species similarity (Jaccard index) with watercourse distance between pairs of sites. The solid lines and 95% confidence intervals (shaded areas) were fitted through linear regression. (**A**) Distance decay for the Oyapock (blue) and Maroni (red) sites sampled with eDNA. (**B**) Distance decay in the Maroni for eDNA (red dots) and capture-based (black triangles) samples considering all captured (or detected) fauna. (**C**) Distance decay in the Maroni for eDNA (red dots) and capture-based (black triangles) samples considering the species captured (or detected) with both capture-based and eDNA methods.

The distance decay of similarity for the sites sampled with gill-nets in the Maroni river, was not significant (p = 0.08, slope= −0.0005, R^2^=0.3), despite a trend of similarity decrease with distance between site pairs (Figure 2**B**). The slope of this trend was significantly less marked than the slope of the similarity decay reported with eDNA data (p<0.001). Restricting the species data to the species detected (or captured) by both sampling methods provided a significant decay of similarity with distance in both eDNA (p<0.001, slope= −0.0009, R^2^=0.6) and capture-based inventories (p=0.022, slope= −0.0006, R^2^=0.47). The decreasing slope of the relationship obtained with the eDNA data was significantly more marked (p<0.001) than the slope obtained with captured based data (Figure 2C).

The comparison of the species spatial ranges along the main channel of the Maroni river provided by the eDNA approach and by gill-net captures revealed a global match of the species distribution between methods (Figure 3). Among the 53 species detected by both methods, 74.5% of the fauna (38 species) was not detected by eDNA downstream from capture-based records. Indeed, distribution ranges were similar between methods for 20 species (39% of the fauna, e.g. *Hoplias aimara*, HAIM; *Eigenmannia virescens*, EVIR; *Brycon falcatus*, BFAL; Figure 3) and for 18 species (35.3% of the fauna) the capture-based method detected the species downstream from the distribution range found using eDNA (*e.g. Moenkhausia georgiae*, MGEO; *Pristobrycon eigenmanni*, SHUM; Figure 3). Only 25.5% of the considered fauna (13 species) were detected using eDNA downstream from the distribution provided by capture-based records (*e.g. Electrophorus electricus*, EELE; *Metaloricaria paucidens*, MPAU; *Geophagus harreri*, GHAR, *Guianacara owroewefi*, GOWR; Figure 3). Furthermore, eDNA and capture-based records retrieved consistent wide distribution ranges along the main channel of the river (*i.e*. species detected in almost all the eDNA and gill-net sites) for 21 species (*e.g. H. aimara*, HAIM; Figure 3). Nevertheless, six species (*G. harreri*, GHAR; *P. barbatus*, PBAR; *Acnodon oligacanthus*, AOLI; *Pseudoplatystoma fasciatum*, PFAS; *E. electricus*, EELE) had a wide distribution range in eDNA records, but not in capture-based records.

**Figure 3:**
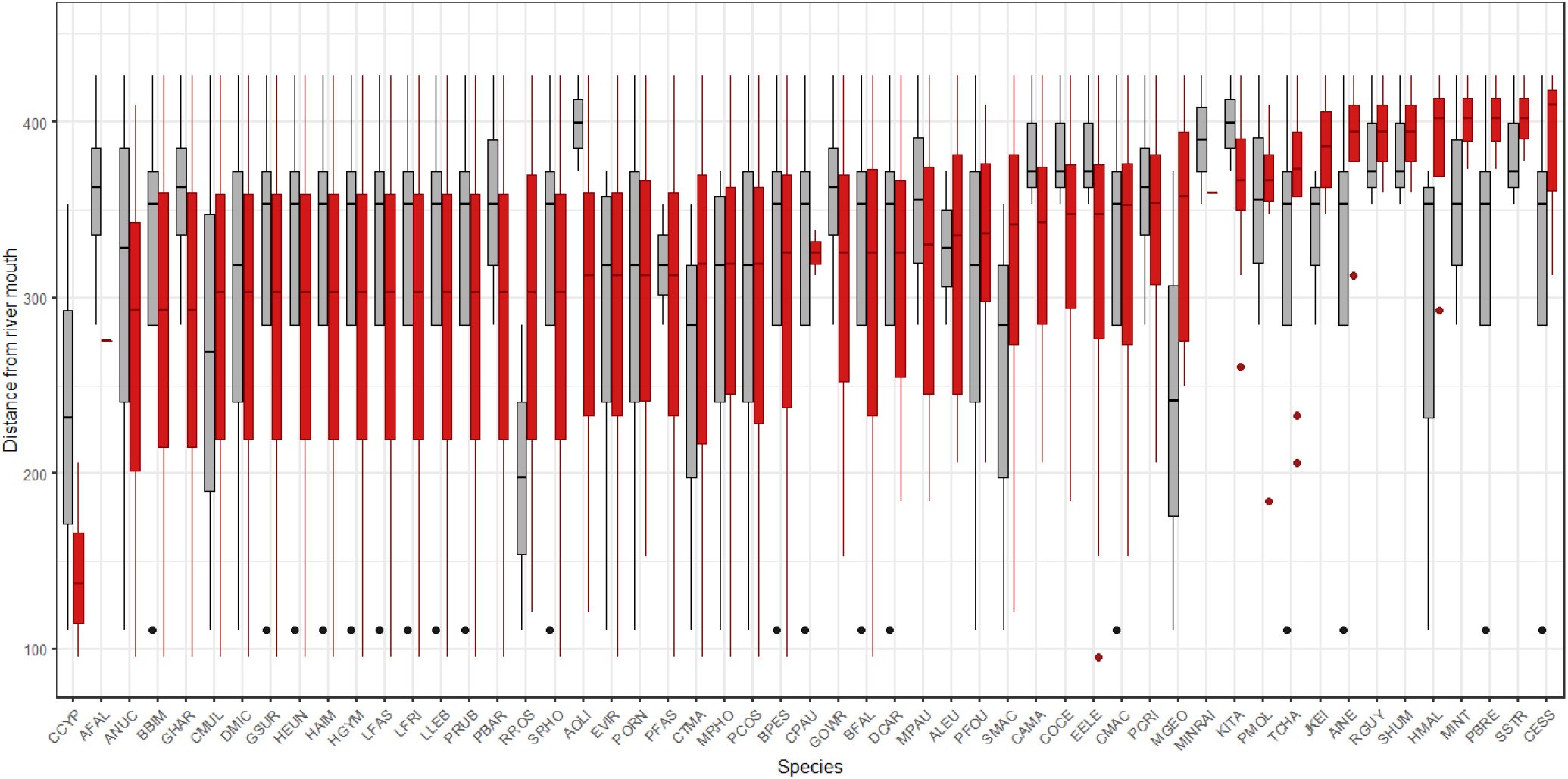
Species distribution ranges along the upstream-downstream gradient of the Maroni river derived from capture-based samples (grey boxplots) and eDNA (red boxplots). Species are ordered according to the mean distance from the river mouth of the sites where they were detected using eDNA (increasing mean from left to right). The species names corresponding to the codes are in Table S1.

## Discussion

As opposed to the ‘conveyor belt’ hypothesis (Deiner et al. 2016) which implies that communities are homogenized across an entire basin because of eDNA transport, our results revealed that eDNA data allowed to retrieve expected spatial patterns of fishes in large tropical rivers at both community and species levels. The decay of similarity between local communities with increasing distance as well as most species distribution ranges were similar to those reported from capture-based samples.

Species similarity in both Maroni and Oyapock rivers was maximal between nearby fish communities, and declined when increasing distances between sampling sites, thus fitting the distance decay theory, a general rule in ecology (Nekola and White 1999). This rule has already been verified in temperate rivers where it reflects the dynamic and heterogeneous architecture of river systems that shape community composition at different spatial scales along the river continuum (Muneepeerakul et al., 2008). However, the distance decay of similarity has rarely been examined in Neotropical aquatic diversity (Araújo et al., 2013). Importantly, the fact that eDNA data was able to reveal distance decay of faunistic similarity patterns necessarily implies a short downstream transport of eDNA. Indeed, according to Nekola and White (1999), the grain size, corresponding to the contiguous area over which a single observation is made (corresponding to the eDNA integration distance in the present study), should be smaller than the whole spatial extent of the study. This allows capturing variation in similarities among comparisons and thus observing distance decay patterns. In our case, this means that the spatial signal of eDNA to detect one fish community need to be smaller than the sampled portion of the river, which is verified for the two rivers, and remains true even when considering only a restricted section of each river. Therefore, the hypothesis of eDNA being transported over long-distances and cumulating downstream was here rejected. Nevertheless, we cannot exclude that the eDNA of some species, probably the most abundant ones, can be transported over long distances as shown by Pont *et al*. (2018) in the Rhone river (France). In the case that this occurs, it would certainly be limited to a few species in our samples as this potential bias did not blurred the significant patterns of distance decay of similarity with geographic distance.

The slopes of similarity decay differed between the two rivers, with a more marked decline in the Maroni river, suggesting a marked decrease of environmental similarities between sites and/or more dispersal constraints with distance in this river compared to the Oyapock (Nekola & White, 1999; Soininen et al., 2007). It can be expected that the decrease of environmental similarities and the increase of dispersal constraints are reinforced by an upstream-downstream gradient of human impacts in the Maroni river, resulting in a higher rate of similarity decay with distance. Indeed, the Maroni river faces six-fold higher deforestation levels than the Oyapock river, (0.37% *versus* 0.06% of catchment area, Gallay *et al*. 2018). Deforestation was mainly caused by gold-mining activities that deeply impact sediment fluxes in the Maroni River (Gallay et al., 2018) and thus strengthen environmental constrains for species (Brosse, Grenouillet, Gevrey, Khazraie, & Tudesque, 2011; Hammond, Gond, Thoisy, Forget, & DeDijn, 2007). This difference between river basins was significantly observed in our eDNA samples, testifying for the relevance of ecological patterns retrieved using eDNA.

Similar patterns of distance decay of similarity in the Maroni river were obtained with eDNA and capture-based data, a pattern which was verified and even reinforced when only the fish species captured by both methods were considered. These results validate, with fish captures, the relevance of the distance decay pattern revealed by eDNA data. Moreover, obtaining similar (and even more marked) distance decay between methods strongly suggests a similar spatial sampling range between methods, and thus reveal that eDNA detection distance is equivalent or even smaller to the size of the reach sampled with gill-nets (*i.e*. from 200 to 500 metres). Interestingly, the negative slope of the eDNA distance decay pattern was significantly more marked than the one retrieved for capture based data. This might either be due to a detection distance for eDNA shorter than the spatial extent of a gill-net sample, or more likely to a higher completeness of eDNA inventories compared to gill-net samples. Indeed, gill-nets are particularly selective in species (Murphy and Willis 1996) and therefore only provide a partial image of the fauna occurring in each site (Cilleros et al., 2019).

Comparing species distribution ranges retrieved from eDNA and capture samples using gill-nets revealed a global match in more than 70% of the species, which corroborates a limited spatial signal of the eDNA detection of the species. More importantly, only 25% of the species were detected using eDNA downstream from their distribution range recorded using capture-based methods. Although this could be considered as species detections due to a downstream drift of eDNA, 10 out of these 13 species have an extended spatial distribution in the Maroni drainage basin and are known from the downstream reaches of the Maroni (Planquette et al. 1996; Keith et al. 2000). The absence of those species in the recent gill-net captures is probably explained by the low capturability of those species due to their morphological and ecological characteristics (Murphy and Willis 1996). For instance, the anguilliform body shape of *E. electricus* (EELE) makes the species difficult to capture with gill-nets, and the sedentary or territorial behaviour of *M. paucidens* (MPAU), *G. harreri* (GHAR), *G. owroewefi* (GOWR) and *K. itany* (KITA) reduces the probability to capture them with passive sampling gears such as gill-nets. Conversely, the 20 species having consistent distributions between methods are species having high capture probability with gill-nets, such as large bodied (*e.g. H. aimara*, HAIM), mobile and gregarious species (*e.g. Brycon falcatus*, BFAL; *Hemiodus unimaculatus*, HEUN), or species with spines and ossified fins that easily tangle in the nets (*e.g. Acuchenipterus nuchalis*, ANUC; *Doras micropoeus* DMIC, *Pimelodus ornatus*, PORN).

## Conclusions

We here demonstrated that fish inventories achieved using eDNA provide an assessment of the fauna at a limited spatial grain, making this biodiversity data appropriate to describe spatial patterns of fish communities and test ecological theories. The scale of eDNA detection certainly does not account for the local habitat of the species that has to be measured over a few square meters. Nevertheless, a detection distance encompassing several hundred of meters appears as a reasonable sample grain size for studying communities or to consider anthropic impacts on fish communities (Jackson, Peres-Neto, & Olden, 2001; Tonn, 1990). Moreover, we show that eDNA can inventory fish fauna at a similar spatial grain than gill-net capture sampling protocols. It might therefore be possible to compare historical and current data on fish faunas, and thus to follow biodiversity alterations experienced by freshwater faunas though current global changes as reported by (Su et al., 2021, p. 202). Moreover, freshwater fish are recognized as one of the most threatened taxa (Hughes, 2021) and the expected species extinction by the end of the century might deeply impact the functional properties of fish assemblages, as well as the services they provide to human societies (Carmona et al., 2021). Under this environmental urgency, eDNA inventories represent a fast and easy to handle fish inventory method providing equivalent and even more precise data than capture based methods.

## Supporting information

Table S1: Species names corresponding to the codes of the Figure 3

## Acknowledgements

This work was funded by DGTM Guyane (project DISTDET), Investissement d’Avenir grant DRIIHM (ANR-11-LABX-0010), through the OHM Oyapock, and the VigiLIFE project. The PAG (Parc Amazonian de Guyane), VigiLIFE and SPYGEN provided field and laboratory support. Investissement d’Avenir projects CEBA (ANR-10-LABX-25-01), TULIP (ANR-10-LABX-0041) and ANR DEBIT (ANR-17-CE02-0007-01) also provided financial support.

## Authors’ contributions

IC and SB conceived the ideas, designed methodology and led the writing of the manuscript; SB, JBD, JM and RV collected the data; IC analyzed the data; AV and TD conducted the laboratory work and bio-informatic analyses.

