## Supplementary material for "Characterizing the spatial signal of environmental DNA in river systems using a community ecology approach": Table S1: Species names corresponding to the codes of the Figure 3

Table S1: Species names corresponding to the codes of the Figure 5.

| Species | Code |
| --- | --- |
| *Acestrorhynchus falcatus* | AFAL |
| *Acnodon oligacanthus* | AOLI |
| *Ageneiosus inermis* | AINE |
| *Ancistrus cf leucostictus* | ALEU |
| *Auchenipterus nuchalis* | ANUC |
| *Bivibranchia bimaculata* | BBIM |
| *Brycon falcatus* | BFAL |
| *Brycon pesu* | BPES |
| *Caenotropus maculosus* | CAMA |
| *Chalceus macrolepidotus* | CMAC |
| *Charax gibbosus* | CPAU |
| *Cichla ocellaris* | COCE |
| *Crenicichla multispinosa* | CMUL |
| *Cteniloricaria platystoma* | CTMA |
| *Curimata cyprinoides* | CCYP |
| *Cynopotamus essequibensis* | CESS |
| *Doras carinatus* | DCAR |
| *Doras micropoeus* | DMIC |
| *Eigenmannia virescens* | EVIR |
| *Electrophorus electricus* | EELE |
| *Geophagus harreri* | GHAR |
| *Geophagus surinamensis* | GSUR |
| *Guianacara owroewefi* | GOWR |
| *Hemiodus unimaculatus* | HEUN |
| *Hoplias aimara* | HAIM |
| *Hoplias malabaricus* | HMAL |
| *Hypostomus gymnorhynchus* | HGYM |
| *Jupiaba keithi* | JKEI |
| *Krobia itanyi* | KITA |
| *Leporinus fasciatus* | LFAS |
| *Leporinus friderici* | LFRI |
| *Leporinus lebaili* | LLEB |
| *Metaloricaria paucidens* | MPAU |
| *Moenkhausia aff intermedia* | MINT |
| *Moenkhausia georgiae* | MGEO |
| *Moenkhausia inrai* | MINRAI |
| *Myloplus rhomboidalis* | MRHO |
| *Pachypops fourcroi* | PFOU |
| *Pimelabditus moli* | PMOL |
| *Pimelodella cristata* | PCRI |
| *Pimelodus ornatus* | PORN |
| *Platydoras costatus* | PCOS |
| *Poptella brevispina* | PBRE |
| *Pristobrycon striolatus* | SSTR |
| *Prochilodus rubrotaeniatus* | PRUB |
| *Pseudancistrus barbatus* | PBAR |
| *Pseudoplatystoma fasciatum* | PFAS |
| *Rhamphichthys rostratus* | RROS |
| *Roeboexodon geryi* | RGUY |
| *Serrasalmus eigenmanni* | SHUM |
| *Serrasalmus rhombeus* | SRHO |
| *Sternopygus macrurus* | SMAC |
| *Tetragonopterus chalceus* | TCHA |
